# Reversible inactivation of different millimeter-scale regions of primate IT results in different patterns of core object recognition deficits

**DOI:** 10.1101/390245

**Authors:** Rishi Rajalingham, James J. DiCarlo

**Affiliations:** McGovern Institute for Brain Research, Massachusetts Institute of Technology; Department of Brain and Cognitive Sciences, Massachusetts Institute of Technology

**Keywords:** object recognition, neural perturbation, inactivation

## Abstract

Extensive research suggests that the inferior temporal (IT) population supports visual object recognition behavior. However, causal evidence for this hypothesis has been equivocal, particularly beyond the specific case of face-selective sub-regions of IT. Here, we directly tested this hypothesis by pharmacologically inactivating individual, millimeter-scale sub-regions of IT while monkeys performed several object discrimination tasks, interleaved trial-by-trial. First, we observed that IT inactivation resulted in reliable contralateral-biased task-selective behavioral deficits. Moreover, inactivating different IT sub-regions resulted in different patterns of task deficits, each predicted by that sub-region’s neuronal object discriminability. Finally, the similarity between different inactivation effects was tightly related to the anatomical distance between corresponding inactivation sites. Taken together, these results provide direct evidence that IT cortex causally supports general core object recognition, and that the underlying IT codes are topographically organized.

## Introduction

Primate core visual object recognition — the ability to rapidly recognize objects in spite of naturally occurring identity-preserving image variability — is thought to rely on the ventral visual stream, a hierarchy of visual cortical areas (DiCarlo et al. 2012). Decades of research suggest that inferior temporal (IT) cortex, the highest level of the ventral stream hierarchy, is a necessary part of the brain’s neural network that underlies core recognition behavior (Logothetis & Sheinberg 1996, Tanaka 1996, Rolls 2000, DiCarlo et al. 2012). For example, it has been shown that parallel linear object discriminants acting on the IT population not only match overall primate behavioral performance (Hung et al. 2005, Zhang et al. 2011) but also predict primate behavioral patterns (Sheinberg & Logothetis 1997, de Beeck et al. 2001, Majaj et al. 2015), showing that IT is a tight neural correlate of primate recognition behavior. Quantitative versions of such experiments have proposed downstream neurally-mechanistic models that successfully link IT population activity to behavior (Majaj et al. 2015) — mechanisms that appear to accurately generalize to all core object recognition tasks. While these experiments are consistent with the hypothesis that IT is a necessary node in the neural network supporting core object recognition behavior, they might also be epiphenomenal (e.g. (Katz et al. 2016, Liu & Pack 2017)). For clarity, we adopt the terminology of Jazayeri & Afraz (2017), whereby causal dependencies link an observed variable (here behavior) to an experimentally controlled variable, in contrast to correlational dependencies, which are associations that we measure and indirectly control, but do not directly control (e.g. associations between neural activity and behavior measured as visual stimuli are experimentally controlled). Thus, to infer a causal dependency between some aspect of IT activity and behavior, it is necessary to directly manipulate IT activity (e.g. via the application of pharmacological agents into IT to silence neurons, etc.) while measuring behavior.

To date, the most successful direct IT manipulations in the context of object recognition have targeted millimeter-scale clusters of face-selective neurons in IT (Afraz et al. 2006, 2015, Moeller et al. 2017, Sadagopan et al. 2017). These studies suggest that neurons in these IT sub-regions are necessary for at least some basic- and subordinate-level face recognition behaviors. Beyond this domain, a notable study by Verhoef et al. (2012) found that manipulation of clusters of 3D-structure preferring neurons in IT influenced the categorization of 3D stimuli as convex or concave. However, results from direct manipulations of IT in general visual object recognition behavior have been equivocal at best. Lesions of IT sometimes suggest the necessity of IT and visual behaviors (Cowey & Gross 1970, Manning 1972, Holmes & Gross 1984, Biederman et al. 1997, Buffalo et al. 2000) but the resulting behavioral deficits are often contradictory (with often no lasting visual deficits) (Dean 1974, Huxlin et al. 2000) and surprisingly modest even for large-scale bilateral removal of IT (e.g. 10-15% drop in performance when complete loss of performance would have been 40%) (Horel et al. 1987, Matsumoto et al. 2016). Thus, it is still unclear if IT is a necessary node in supporting general core object recognition behavior. Moreover, even if IT cortex is indeed necessary for all core object recognition tasks, it is unclear if that assumed causal role is spatially organized. For example, the current literature on monkey IT is consistent with the hypothesis that every square millimeter of IT cortex outside of the fMRI-defined face patches is equally involved in all (non-face) object discriminations, with some authors implicitly arguing for that hypothesis (Tsao & Livingstone 2008, Kanwisher 2010).

To investigate these open questions, we here reversibly inactivated neurons in individual, arbitrarily sampled millimeter-scale regions of IT via local injection of muscimol while monkeys performed a battery of pairwise core object discrimination tasks, interleaved trial-by-trial. This paradigm allowed us not only to directly test the aforementioned IT-to-behavior linking hypotheses (Majaj et al. 2015), but also to characterize the causal role of each inactivation IT site via a *pattern* of deficits over object recognition tasks.

Our results show that inactivation of even single, millimeter-scale regions of IT resulted in reliable contralateral-biased behavioral deficits. Interestingly, these deficits were highly selective over core object recognition tasks — inactivating a small region of IT produced deficits in only a subset of such tasks, and inactivating different such regions resulted in different patterns of object recognition deficits. Furthermore, the effect of inactivation was topographically organized in that the pattern of behavioral deficit (i.e. the pattern over tasks) was most similar at anatomically neighboring injection sites. We also found that each pattern of task deficit was well predicted by the object discriminability of the local region’s neuronal activity. Taken together, these results demonstrate the necessity of IT cortex for a wide range of general core object recognition behaviors, and reveal that — even outside of face patches — IT cortex has behaviorally-critical topographic organization of visual features. These findings are consistent with and suggested by prior physiology work (Wang et al. 1998, Tsunoda et al. 2001, Kreiman et al. 2006), but, to our knowledge, this is the first demonstration of a topographically organized causal role of IT in general core object recognition.

## Results

Our primary goal was to ask if IT causally supports object recognition, and whether any such causal role is functionally specific at the millimeter-scale, as schematized in Figure 1A. To do this, we reversibly inactivated individual, arbitrarily sampled millimeter-scale regions of IT via injection of muscimol while monkeys performed a battery of pairwise core object discrimination tasks. Figure 1B shows the behavioral paradigm used for testing monkeys’ core object recognition behavior. In this work, the battery consisted of 6 (Monkeys 1, 2) or 10 (Monkey 2 only) pairwise core object discrimination tasks between five objects, interleaved trial-by-trial (see Figure 1 A for task list, Figure S1 for control behavior). To enforce true object recognition (rather than image matching), stimuli consisted of naturalistic synthetic images of 3D objects rendered under high view-uncertainty (see S1 for example images), and the monkey subjects were required to generalize to new images in each task (as we have previously shown they readily do (Rajalingham et al. 2015)). Figure 2A shows the behavioral data for an example inactivation experiment in Monkey 1, for each of six pairwise discrimination tasks. Each panel shows the relative behavioral performance (mean ± SEM, obtained by bootstrap resampling over trials) for a given pairwise task, for each of three consecutive behavioral sessions (pre-inactivation control, inactivation, and post-inactivation control; see Methods). Performance on each task is shown relative to the average performance on that task over the pre- and post-control sessions, which we use as a measure of control behavior on each task (see Methods); the dark and light shaded areas correspond to one and two SEM of this measure (computed over trials), respectively. We observed a strong and significant deficit for some tasks (i.e. chair vs. dog, chair vs. plane, and dog vs. bear) but not others (elephant vs. bear, dog vs. elephant). The resulting pattern of behavioral deficits (i.e. the deficit pattern over tasks) for this one example inactivation site in IT is shown in Figure 2B, with the corresponding anatomical location shown in the inset. For this volume of injection, we expect strong neural suppression in a volume ~ 2.5mm in diameter centered at the injection site (Arikan et al. 2002). Figure 2C shows the pattern of behavioral deficits for eight more example inactivation sites in IT out of the 25 sites we tested from both monkeys (Monkey M, P in the first and second row, respectively). We qualitatively observe that inactivating each local IT region resulted in reliable task-specific behavioral deficits, and that the pattern is not the same for all IT regions. Together, these results suggest that inactivating different millimeter-scale regions of primate IT results in deficits in different core object recognition tasks (i.e. different patterns of deficits). This inference is directly and quantitatively tested in the following analyses.

**Fig. 1.**
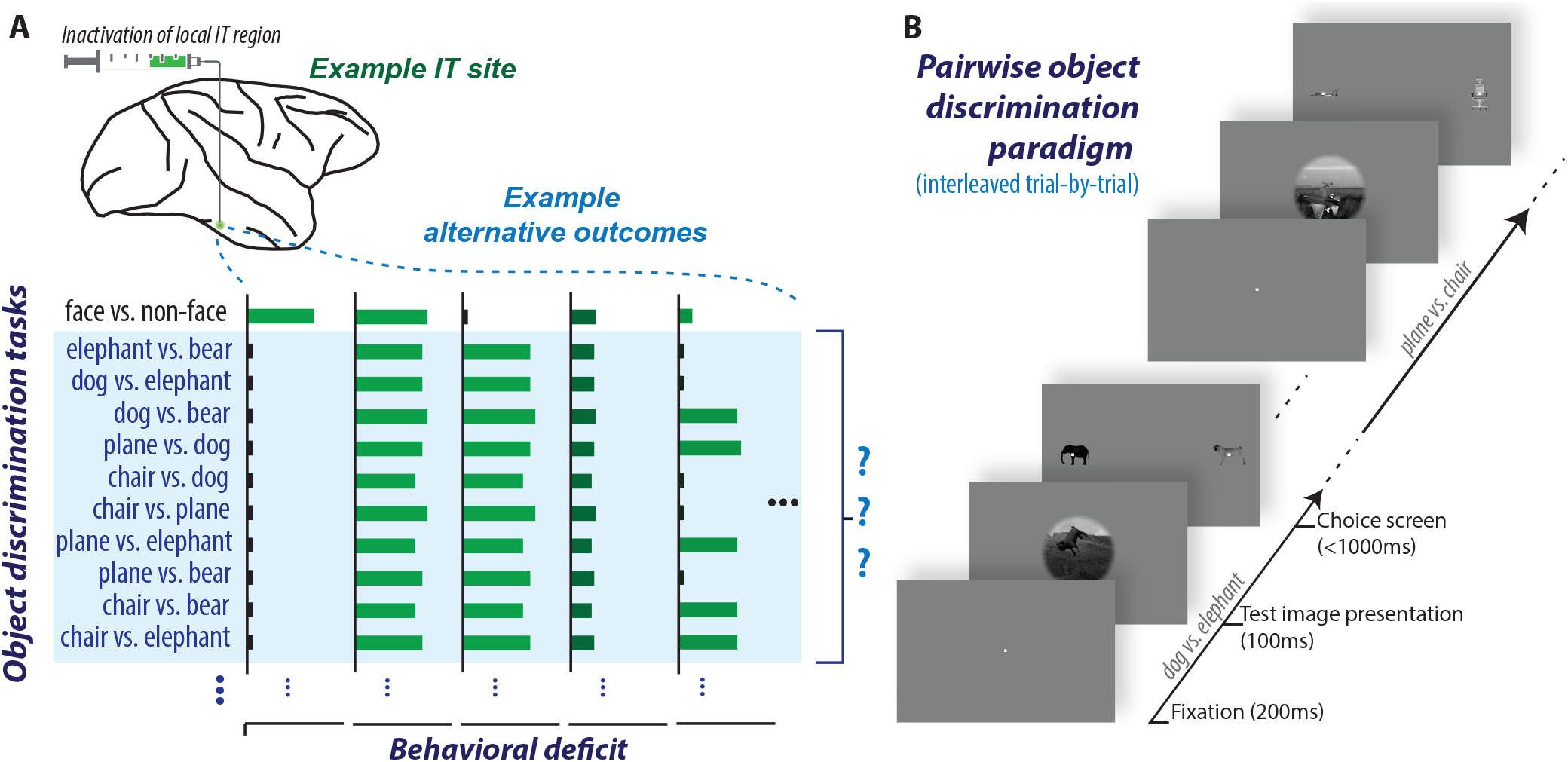
(a) Schematic of experiment. It is still unclear if IT is necessary for general core object recognition behavior, and moreover if any such causal role is functionally specific at the millimeter-scale. To investigate this, we reversibly inactivated individual arbitrarily-sampled millimeter-scale regions of IT via local injection of muscimol while monkeys performed a battery of pairwise core object discrimination tasks (listed, highlighted in blue), interleaved trial-by-trial (see b). Bar plots outline alternative possible outcomes corresponding to different patterns of behavioral deficits from such inactivations, varying in task-selectivity from highly specialized (far left panel, exhibiting deficits only for face vs. non-face discriminations), to largely uniform (middle three panels, exhibiting equal deficits on all discrimination tasks, or all non-face discrimination tasks), to relatively task-selective (far right panel, exhibiting deficits on some but not all discrimination tasks). (b) Behavioral paradigm. Each trial was initiated when the monkey acquired and held its gaze on a central fixation point for 200ms, after which a test image (8×8 degrees of visual angle) appeared at the center of gaze for 100ms. After extinction of the test image, two choice images, each displaying a single object in a canonical view with no background, were immediately shown to the left and right. One of these two objects was always the same as the object that generated the test image (i.e. the correct choice), and its location (left or right) was randomly chosen on each trial. The monkey was allowed to freely view the choice images for up to 1000ms, and indicated its final choice by holding fixation over the selected image for 700ms. A juice reward was delivered immediately after each correct trial. Note that we refer to each pairwise object discrimination (averaged over all test images) as a “discrimination task” and the trials for all such tasks (6-10 tasks, see Methods) were pseudo-randomly interleaved trial-by-trial.

**Fig. 2.**
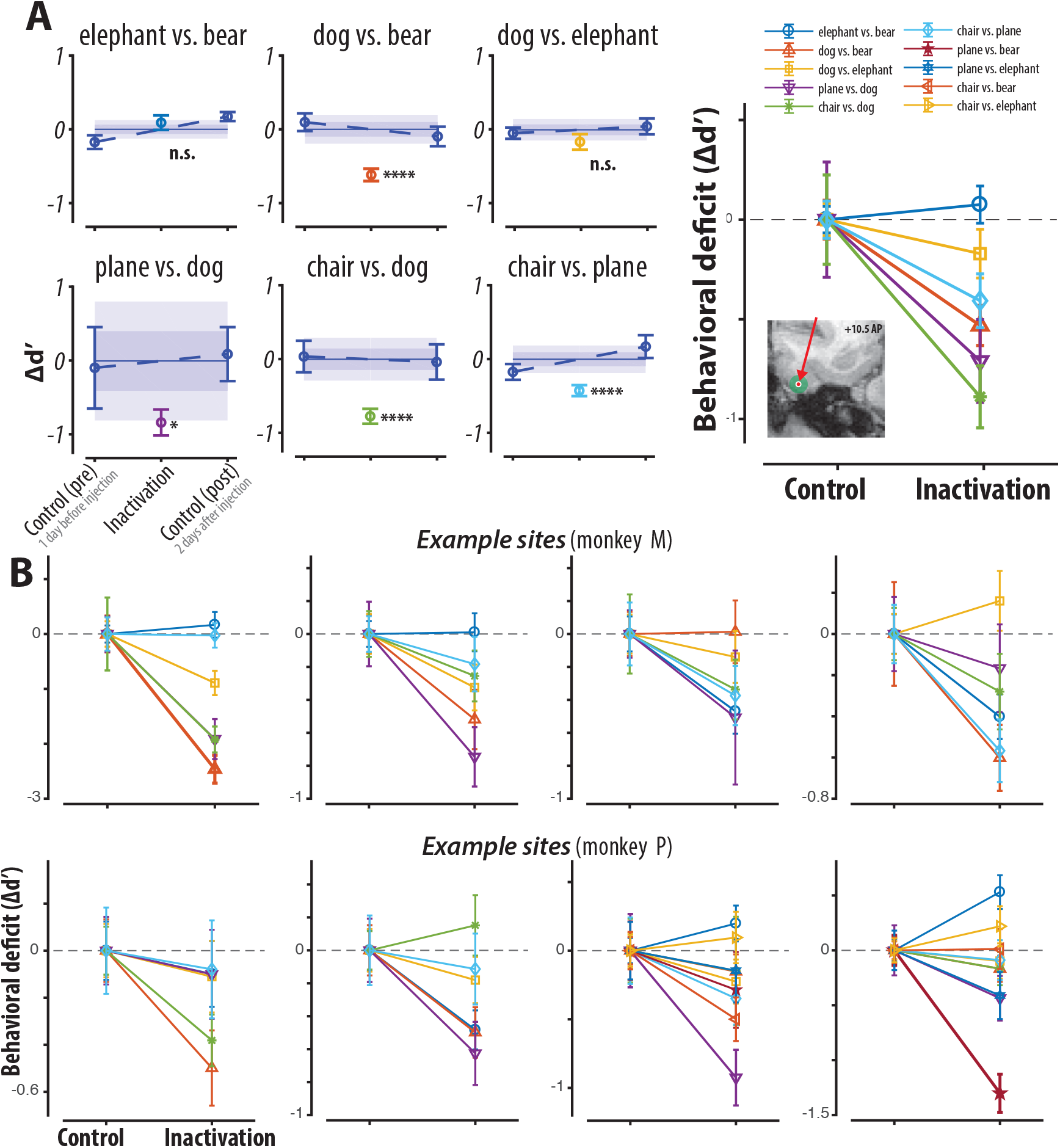
(a) Example inactivation experiment. (left) Behavioral performance (mean) for each of six tasks over the three condition (pre-control, inactivation, and post-control; see Methods). Data are shown as behavioral performance relative the average of pre- and post-control performances (see Methods) (bars show SEM obtained by bootstrap resampling over trials). The location of the injection site (“inactivation” condition) for this experiment is shown in the right panel. The dark and light shaded areas correspond to one and two SEM respectively of this control. For this site, we observed a strong and significant deficit for some tasks (chair vs. dog, chair vs. plane, and dog vs. bear) but not others (elephant vs. bear, dog vs. elephant). (right panel) The data on the left are summarized relative to the average control performance (mean ± SEM over trials). (b) Eight more example IT inactivation experiments using the same format as (a, right), out of the 25 sites we tested. Note that behavioral performance on one or more object discrimination tasks is typically reduced by inactivation of each IT location, the specific task(s) affected are different at different IT locations, and no IT location affected all the tasks.

### Summary of behavioral deficits

Figure 3A shows the behavioral deficits for all inactivation sites and all tasks in both monkeys as a scatter of control performance versus inactivation performance (n=25 sites, n=182 tasks × sites). Considering all the tasks together, we observed a significant decrease in performance (i.e. inactivation lower than control), corresponding to the predominance of points under the unity line in Figure 3; on average, this amounted to a global deficit of *μ_δ_* = –0.2 ± 0.02 in units of d’ (*p* = 1.23 * 10^−16^, one-tailed exact test; see Figure 3B, red bar under *global deficit*). Consistent with the known lateralization of IT (Op De Beeck & Vogels 2000), this deficit was more pronounced for images in which the center of the target object was contralateral to the injection hemisphere (*μ_δ_* = –0.26 ± 0.03, *p* = 1.28*e* – 16) than for images with ipsilateral object centers (*μ_δ_* = –0.17 ± 0.03, *p* = 3.82 * 10^−12^) and this difference was significant (*p* = 0.0128, one-tailed exact test; ipsi vs. contra). Note that all images were presented foveally (spanning –4° to 4° of both azimuth and elevation, and average object size was ~ 3.5°).

**Fig. 3.**
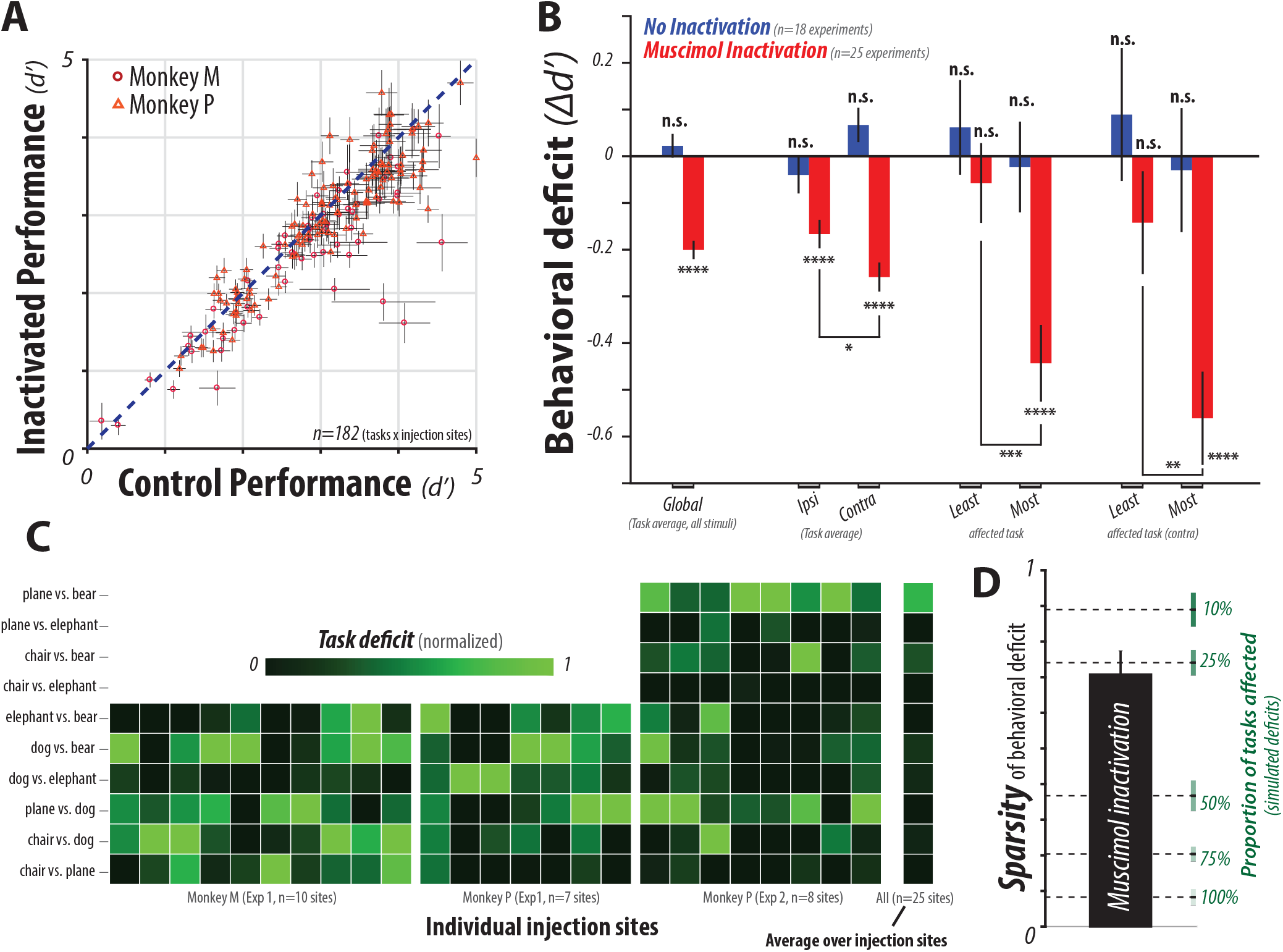
(a) Behavioral deficits for all IT inactivation sites and all tasks in both monkeys as a scatter of control performance and inactivation performance. Note the on-average decrease in performance corresponding to predominance of points under the unity line (dashed line). (b) Summary of behavioral deficits when grouping the tasks and task images in different ways. Red bars show the magnitude of inactivation deficit (relative to control) for each grouping. From left to right, these groupings are: all images and all tasks (“Global”), ipsilateral/contralateral object images for all tasks (“Ipsi” and “Contra”), the least/most affected task at each site (“Least” and “Most”) selected on held out data, contralateral object images for the least/most affected task at each site (“Most Contra”) selected on held out data. Blue bars correspond to otherwise identical experiments but without muscimol inactivation (control experiments). (c) The heat map shows the task deficits for each of the 25 inactivation sites, with brighter colors corresponding to larger relative task deficits, highlighting that inactivation of each IT site resulted in a different, relatively sparse, pattern of behavioral deficit. The average deficit pattern over all inactivation sites (right column) is largely uniform, suggesting that IT as a whole is approximately equally involved in each discrimination task. (d) The black bar shows the sparsity (see Methods) of the behavioral task deficits, over all sites (mean ± SEM over sites). To provide calibration, green lines show the sparsity values that occur under simulations in which we varied the proportion of truly affected tasks and used identical sampling noise as our data; see Methods). Together, these results suggest that inactivation of a single 2.5 mm diameter region of IT affects 25% of core object discrimination tasks, on average.

Next we asked whether the inactivation deficits were task-specific. To examine this, we compared the magnitude of behavioral deficits between the least-affected and most-affected tasks, for each inactivation site. Crucially, to avoid any selection bias, these tasks were selected from held-out data: we split our data into two disjoint halves of trials, selected the least/most affected tasks per inactivation site from one split-half, and examined the corresponding deficits on these selected tasks in the second split-half (thus the expected value of the difference in deficits between the most and least affected task is zero under the null hypothesis; see Methods). Using this procedure, we observed a large significant behavioral deficit for the most affected task (*μ_δ_* = –0.44 ± 0.08, *p* = 2.79 * 10^−16^), but not for the least-affected task (*μ_δ_* = –0.06 ± 0.08, *p* = 0.27), and the difference was significant (*p* = 3.99 * 10^−4^, see Figure 3B). Finally, we observed even larger task-selective deficits when restricting to contralateral objects (as described above), with a similar significant difference between the most and least affected tasks (*μ_δ_* = –0.56 ± 0.1, –0.14 ± 0.11 for most and least affected tasks respectively; *p* = 1.44 * 10^−3^).

For each of the analyzed conditions, we observed no significant behavioral deficits on otherwise identical experiments without muscimol inactivation (*p* > 0.05; Figure 3B, blue bars). Furthermore, the patterns of deficits across these analyzed conditions were similar for both animals (Figure S2). In summary, inactivation of local regions of IT resulted in highly reliable behavioral deficits, which were selective over visual space (i.e. contralateral-biased) and selective over different core object recogntion tasks.

### Task-selectivity of deficits

Focusing on “contralateral stimuli” (i.e. images in which the center of the target object was contralateral to the injection hemisphere), Figure 3C shows the task deficit patterns for each of the 25 inactivation sites as a heat map. Each column corresponds to the deficit pattern over tasks from inactivating an individual IT site, normalized to a fixed color scale ([0,1]); brighter colors correspond to larger relative task deficits. Consistent with the inferred task-selectivity from Figure 3B, we observe that each inactivation resulted in a non-uniform behavioral deficit pattern. This non-uniformity was quantified via a sparsity index (*SI*, see Methods) which has a value of 0 for perfectly uniform deficit patterns (i.e. where each IT sub-region is equally necessary for all tasks), and a value of 1 for a perfectly task-specialized (or “one-hot”) deficit pattern (i.e. where each sub-region is necessary for just one of the tested tasks). We observed that inactivation of local regions in IT led to highly non-uniform deficit patterns, on average (*SI*(*δ*) = 0.71 ± 0.05; mean±SEM over sites, see Figure 3D).

To ground this empirical *SI* value, we estimated the corresponding SI distributions for different simulated behavioral deficit patterns with varying degrees of non-uniformity across tasks. These simulated deficit patterns were obtained via random permutations of our data, varying only the proportion of affected tasks (see Methods), in an effort to best match all other sources of variability (e.g. trial and task sampling variability) for direct comparison with the empirically observed *SI*. Figure 3D shows the SI distributions expected from behavioral deficits of varying degrees of nonuniformity (i.e. with 10%, 25%, …, 100% of tasks affected). We observe that the empirically observed task-selectivity is significantly greater than expected from a uniform deficit (*p* = 2.42 * 10^−16^; relative to simulated 100%-affected, i.e. uniform) but significantly less than expected from a highly sparse deficit pattern (*p* = 5.28 * 10^−3^; relative to simulated 10%-affected). Indeed, the observed SI estimates correspond to simulation of deficits on ~ 25% of tested tasks.

Importantly, this non-uniformity does not simply reflect nonuniformity in the behavioral difficulty across tasks. Indeed, normalizing each deficit pattern by the behavioral difficulty pattern resulted in normalized deficit patterns that were not significantly correlated with task difficulty (*r* = 0.06, *p* = 0.39), and significantly non-uniform as quantified by sparsity (*SI*(*δ_n_*) = 0.74± 0.06; *p* = 2.21 * 10^−6^, relative to simulated uniform). This is also clear from Figure 3C, which shows that inactivation of different sites led to different deficit weight patterns (left panel). Accordingly, the deficits were relatively evenly distributed over the tasks, as reflected by the uniformity of the the average deficit pattern over all sites (3C, right-most bar). Together, these results indicate that the nonuniformity of task deficits is not tied to specific tasks.

### Tissue selectivity of deficits

Inactivating different anatomical regions of IT resulted in different patterns of task deficits. To directly test this tissue selectivity, we compared the inactivation deficit patterns between pairs of IT sites. Pairwise deficit pattern similarity was quantified using a noise-adjusted correlation (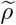, see Methods). We considered all pairs of inactivation sites, measured within the same animal and image-set, where the inactivation deficit patterns of both sites had split-half internal reliability greater than a threshold *θ* (*n* = 62 pairs for *θ* = 0.1, but results did not significantly depend on the choice of the threshold *θ*). We measured the dependence of pairwise deficit similarity on the anatomical distance between the inactivation sites, where anatomical distance (d) was computed as the Euclidean distance between the injection site locations estimated via high-resolution micro-focal stereo x-ray reconstruction (see Methods). We first observed that inactivation deficits are highly replicable across experiments: the noise-adjusted correlation between behavioral deficit patterns of neighboring inactivation sites was near ceiling (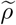 = 0.92 ± 0.03 for *d* < 1mm, mean ± SEM over site pairs; Figure 4). We further observe that this similarity between the inactivation deficits of two injection sites was monotonically related to the anatomical distance between (Figure 4). A simple exponential decay model (half-max-full-width *HMFW* = 3.29 ± 1.19mm) significantly explained this relationship (*R*^2^ = 0.36 ± 0.12, *p* = 8.04 * 10^−4^). We verified that this model correlation is not expected by chance by fitting the model on randomly shuffled data (*R*^2^ = 0.00 ± 0.13, *p* = 0.50). Note that this *HMFW* estimate is approximately consistent with a combined effect of the known spatial spread of muscimol in the cortical tissue and previously reported columnar organization of IT at a sub-millimeter-scale. Together, these results suggest that behavioral deficits are tissue-specific, i.e. the effect of inactivation is different for different inactivation sites, and most similar at anatomically neighboring injection sites.

**Fig. 4.**
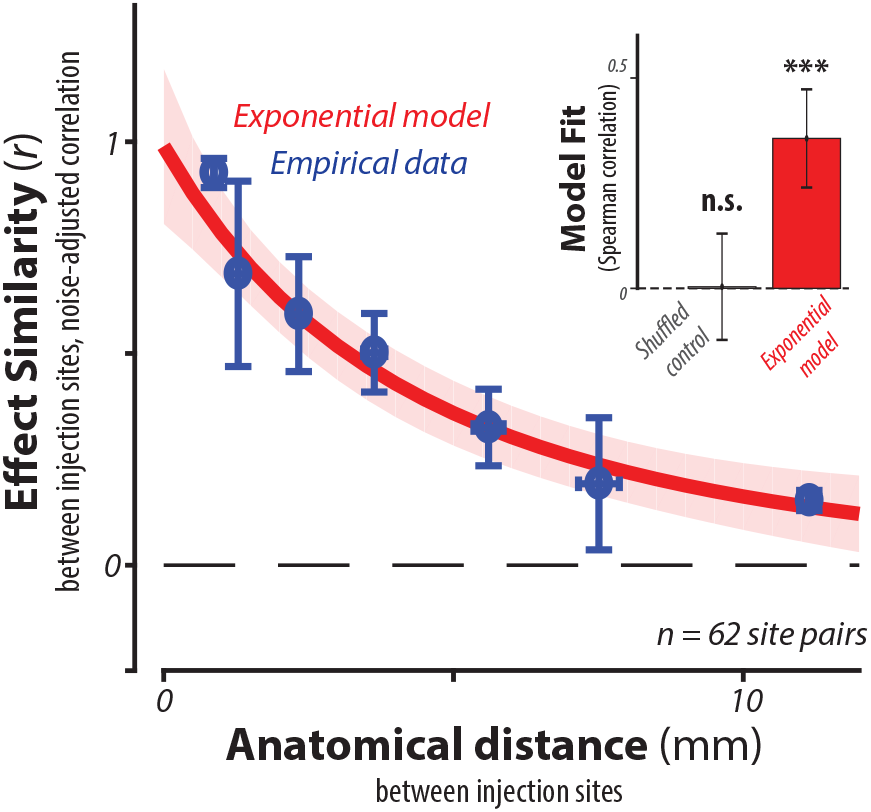
(a) Topographical organization. The main panel plots the similarity in behavioral deficit patterns between pairs of IT injection sites (quantified as noise-adjusted correlation, y-axis) as a function of the anatomical distance between each pair of sites (x-axis). Empirical data are shown as the mean (± SEM) of all pairs of sites in logarithmically-spaced bins of tissue distance (blue points). Note that the pattern of inactivation-induced behavioral effects is highly replicable in that we observe very high correlation of effects for repeated experiments at or very near the originally tested site (near 0 on the x-axis). The similarity between any two inactivation deficits was monotonically related to their anatomical distance, and a simple exponential model significantly explained this relationship (see inset).

### Neurally-mechanistic models that link IT activity to behavior

Given the observed tissue specificity, we asked to what extent the observed behavioral deficits could be predicted by the neuronal activity patterns in the inactivated sub-regions (e.g. prior to inactivation). The central panel in Figure 5A shows the location of an example muscimol inactivation site and local electrophysiology sites, co-registered using stereo micro-focal x-ray reconstruction, and overlaid on a coronal MRI slice. For this example site in IT, we recorded the activity of eight multi-unit sites (shown as cyan discs) in close proximity to the injection site (shown as red disc). Multi-unit activity was recorded in response to the same images as those used in behavioral testing, in a passive viewing paradigm (see Methods). Each sub-panel shows a multi-unit site’s stimulus-locked firing rate responses for each of the five objects, averaged over images. We note that neuronal sites, while heterogeneous, each exhibit reliable object preferences. Based on local neuronal responses such as this, we constructed and tested a number of linking (aka “decoder”) models, each of which maps the local IT spiking response patterns to a predicted behavioral deficit.

**Fig. 5.**
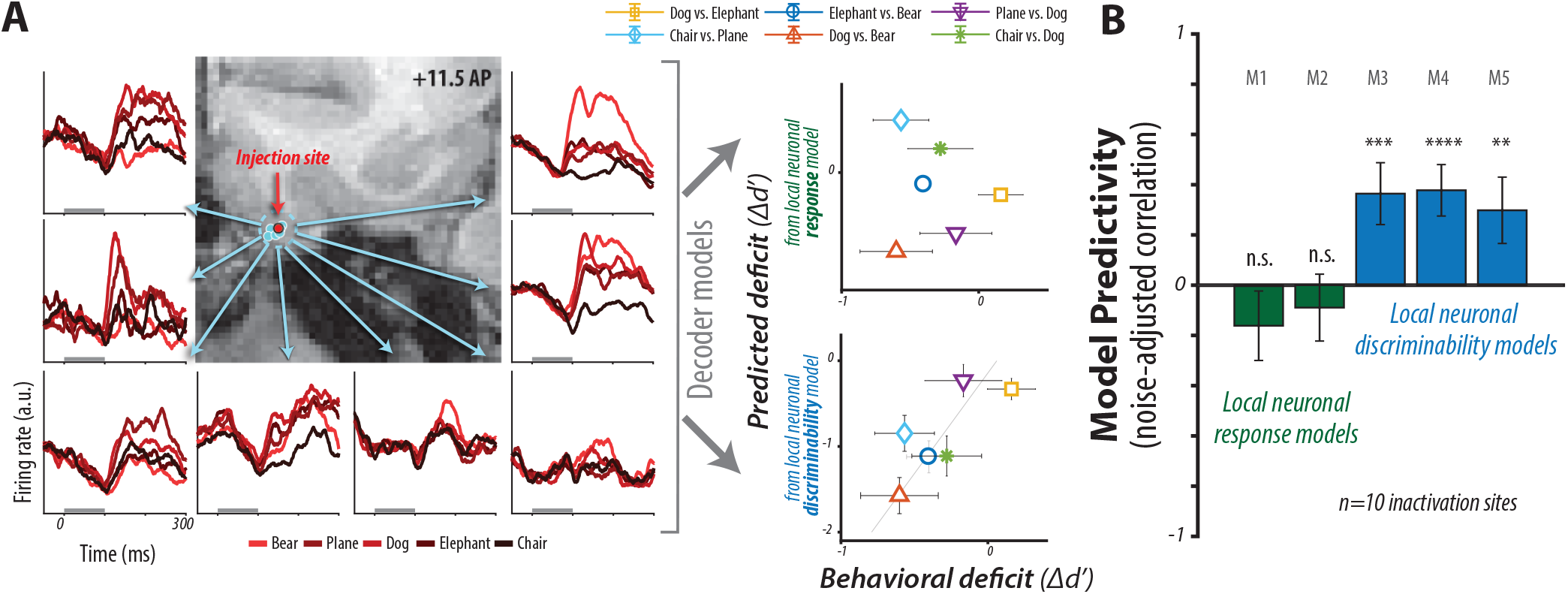
Relationship of IT spiking responses to patterns of behavioral deficit. (a) Left: Multi-unit spiking activity recorded serially (prior to inactivation) with a single microelectrode for eight sites sampled within an example IT sub-region. The recording locations (each determined via stereo, micro-focal X-ray; see Methods) are here plotted projected into the plane of a single MRI slice containing the center of the IT inactivated region. Each inset panel shows the spiking activity response to each of five objects aligned to stimulus onset; each line is the mean activity, averaged over all images of each object and all repetitions (40 images/object, ~ 10 repetitions/image). Gray bar shows image presentation time (100 ms). Neuronal sites, while heterogeneous, each exhibit object preferences, even when averaging over images. Right: To determine whether the observed behavioral deficits are predicted by local neuronal activity, we constructed and tested several decoder models that transform IT response patterns into predictions of behavioral deficits resulting from inactivation (see Methods). The predictions of two of these models (upper and lower scatter plot) are compared with the measured behavioral deficits for this example IT inactivation site. Note that larger deficits correspond to more negative values of Δ*d*’ (lower left corner of each scatter plot). (B) The average predictive power of each of five tested decoder model is shown as the noise-adjusted correlation between predicted and actual behavioral deficits, for all relevant sites (i.e. where we had both the local spiking responses (as in a) and the pattern of behavioral deficits measure on the same set of images). Each bar corresponds to a specific decoding model (models M1-M5, see Methods). All local neuronal discriminability models (blue) were clearly better than the local neuronal response models (green).

Figure 5B shows the predictions from two example linking models. The local neural response models predict large deficits for tasks with images that produce the largest response from the local neuronal sites. The local neural discriminability models predict large deficits for tasks for which the local neural spiking activity was most discriminative, as measured by a linear classifier. We qualitatively observe that the discriminability models better capture the observed behavioral deficit patterns than the response models, for this example inactivation site. This is quantified in Figure 5C as a noise-adjusted correlation between predicted and actual behavioral deficits, over all inactivation sites with local neuronal recordings (*n* = 10 sites for *d* < 2mm). All discriminability models significantly predict the inactivation deficits (*p* < 0.001), while the response models failed to do so (*p* > 0.05). In summary, inactivation of millimeter-scale regions of IT results in behavioral deficits that are predicted by the local neuronal discriminability.

## Discussion

In this work, we sought to investigate *if* and *how* neural activity in IT causally supports core object recognition behavior. Specifically, our goals were to 1) directly test the hypothesis that IT is a necessary node in the brain’s neural network that underlies potentially all core object recognition discrimination behavior (tasks), and 2) to ask if any such causal role is functionally organized over the cortical tissue. To this end, we reversibly inactivated individual, arbitrarily sampled millimeter-scale regions of IT while monkeys performed a battery of object discrimination tasks. Our first contribution is to provide direct causal evidence for the role of IT in core object recognition, which was largely lacking, especially beyond the specific case of face-selective subregions of IT. Moreover, our results revealed that the causal role of IT in object recognition has topographic organization at the millimeter-scale and is predicted by local neuronal discriminability. Together, these advances solidify the previously-presumed causal role of IT cortex in core object recognition, and could be used to distinguish among alternative neurally-mechanistic (i.e. neural network) models of the ventral stream and its role in core object recognition behavior, as outlined below.

### The hypothesized role of IT cortex in core object recognition behavior

We here define the decoding hypothesis (aka linking hypothesis (Brindley 1960)) that motivated the present study, and alternatives to that hypothesis. First, we hypothesize that IT cortex is a necessary node in the brain’s neural network that underlies core recognition behavior (*Prediction 1*). Stated in other words, our hypothesis is that core object recognition behavior causally depends on the firing of neurons in IT cortex, and without those spikes, core object recognition behavior would be at chance (DiCarlo et al. 2012, Majaj et al. 2015). Importantly, core object recognition behavior is not a single “task”, but is a domain of many possible tasks — including at least hundreds of pairwise object discrimination tasks in monkeys (Rajalingham et al. 2015, 2018). Thus, based on prior IT recording work (Majaj et al. 2015), our decoding hypothesis is more specific: each IT neuron is a necessary part of multiple such tasks (*Prediction 2*), which is contrasted with the alternative possibility that all non-face-selective IT neurons are necessary for *all* object tasks. Third, our decoding hypothesis is that single IT neurons that carry information that might *potentially* support each task, are indeed *necessary* for each such task, and they are necessary *regardless of their physical location in IT* (*Prediction 3*). This hypothesis is implicitly stated in (Majaj et al. 2015) and explicitly discussed in (Afraz et al. 2015). However, because prior work (Tanaka 1996, Kreiman et al. 2006, Sato et al. 2008) showed that IT neurons with similar object feature and image preferences tend to be clustered at millimeter-scale, our decoding hypothesis (above) predicts that each mm-scale IT sub-region is an enrichment of neurons that are necessary nodes in some object discrimination tasks (again, more than one task). This is contrasted with the alternative possibility that each sub-region of IT is equally involved in all object discrimination tasks. We note that all of these assumptions (here collectively called our “decoding hypothesis”) and the resultant predictions (*Predictions 1-3*) were in place prior to our undertaking of this study, and indeed, were the motivation of this study.

### Direct causal evidence for the role of IT in core object recognition

While we cannot yet test all the mechanistic aspects of this decoding hypothesis (above), we can test some it most basic predictions — to our knowledge, these tests had not yet been done. To carry out these tests, we adopt the terminology of Jazayeri & Afraz (2017), whereby “causal” dependencies can be inferred by correlating a dependent variable to an experimentally controlled variable, in contrast to correlational dependencies which are associations between variables that we measure and may indirectly control, but we do not directly control. Thus, to infer a causal link between IT activity and behavior, it is necessary to directly manipulate activity in IT (e.g. via the application of pharmacological agents into IT to silence neurons, etc.) while measuring behavior. Related correlational dependencies (e.g. via direct manipulation of visual input to the retinae while measuring variations from both IT activity and behavior) are consistent with our causal decoding hypothesis (outlined above) but could also reflect epiphenomenal mechanisms; i.e. correlation does not imply causation. Recently, research in other behavioral domains has exposed divergences between correlational and causal dependencies (Katz et al. 2016, Liu & Pack 2017), highlighting the need to directly test causal dependencies.

With respect to *Prediction 1* of our stated decoding hypothesis (that IT is necessary for core object recognition), decades of neurophysiological and neuropsychological research suggest that activity in IT cortex is a good neural correlate of primate object recognition behavior (Logothetis & Sheinberg 1996, Tanaka 1996, Rolls 2000, DiCarlo et al. 2012): individual neurons in IT cortex are selective to complex visual features in images, and exhibit remarkable tolerance to changes in viewing parameters (Kobatake & Tanaka 1994, Ito et al. 1995, Logothetis et al. 1995, Booth & Rolls 1998, Rust & DiCarlo 2010), and the population of neurons in IT not only matches overall primate behavioral performance (Hung et al. 2005, Zhang et al. 2011) but also reliably predicts the behavioral error patterns (Majaj et al. 2015). Taken together, these results are consistent with our decoding hypothesis, but could also reflect epiphenomenal mechanisms. To this end, our first major contribution in this work is to provide direct causal evidence for the role of IT in core object recognition behavior. Prior to this, causal evidence for the role of IT in core object recognition has been both scarce and equivocal, especially beyond the specific case of face-selective regions in IT. Lesions of IT suggest a coarse causal link between this area and visual behaviors (Cowey & Gross 1970, Manning 1972, Holmes & Gross 1984, Weiskrantz & Saunders 1984, Buffalo et al. 1998, Huxlin et al. 2000, Matsumoto et al. 2016) but the resulting behavioral deficits are often contradictory (Dean 1974, Huxlin et al. 2000) and at best modest (Horel et al. 1987, Matsumoto et al. 2016). For example, recent work showed that near complete ablation of IT (bilateral removal of anterior IT) resulted in only mild (10-15%) deficits in object categorization (Matsumoto et al. 2016). It is unclear to what extent these modest behavioral deficits can be explained by limitations of the methodologies and the behavioral assays, which may not be robust to alternative (potentially compensatory) strategies. A handful of studies have reported using focal reversible neural perturbation tools (e.g. electrical, pharmacological, and optogenetic perturbation) to test the stated decoding hypothesis, but all exclusively targeted spatial clusters of face-selective neurons in IT, testing the causal role of these regions in basic- and subordinate-level face recognition behaviors (Afraz et al. 2006, 2015, Moeller et al. 2017, Sadagopan et al. 2017), with one notable exception (Verhoef et al. 2012). Thus, our results provide the most systematic direct causal evidence for the general decoding hypothesis (i.e. *Prediction 1)* outlined above.

### The causal role of IT in core object recognition is topographically organized

With respect to *Prediction 2* of our stated decoding hypothesis (that each mm-scale IT subregion is necessary for several, but not all, object discrimination tasks), our second major contribution in this work is to provide direct evidence for a task-selective causal role of IT in core object recognition at the millimeter-scale. Prior to this, all existing studies have exclusively targeted specific spatial clusters of face-selective neurons in IT, testing the causal role of these regions in basic- and subordinate-level face recognition behaviors (Afraz et al. 2006, 2015, Moeller et al. 2017, Sadagopan et al. 2017). While faces are an especially behaviorally relevant stimulus class for primates (Tsao & Livingstone 2008), the experimental bias towards face-preferring (on average) spatial clusters in IT is likely related to the spatial resolution limitations of current neural perturbation tools, which operate on groups of spatially contiguous neurons at approximately millimeter-scale. Given this limitation, the known millimeter-scale spatial clusters of face selective regions in IT (Tsao et al. 2003, 2006, Tsao & Livingstone 2008) form an intuitively optimal candidate for testing causal dependencies related to our decoding hypothesis. We note that similar spatial clustering of response selectivity has been reported for a small number of other image groupings besides faces, such as color, places, and bodies (Conway et al. 2007, Kornblith et al. 2013, Lafer-Sousa & Conway 2013, Verhoef et al. 2015, Popivanov et al. 2012). Importantly, the topographic organization of neurons in IT is largely unknown and assumed by many to be functionally random and nonspecific beyond these discrete clusters. To support a general inference, we here tested arbitrary sampled millimeter-scale regions of ventral IT, rather than functionally target inactivation sites. Interestingly, we found that inactivation of different regions in ventral IT led to different task-specific deficits, suggesting some functional specificity for arbitrarily sampled millimeter-scale regions. Indeed, our data suggest that each millimeter-scale region in IT is causally involved in a relatively small proportion (~ 25%) of object recognition tasks, and that anatomically neighboring regions are similar in this regard. This topographical organization is consistent with previously reported sub-millimeter scale columnar organization of neurons in IT (Fujita et al. 1992, Tanaka 1996, Wang et al. 1996, 1998, Kreiman et al. 2006). We speculate that this topographic organization could reflect a general principle of global cortical layout, whereby neuronal selectivities are developed in the face of metabolic constraints (e.g. minimization of connection wiring length (Chklovskii et al. 2002)).

### The causal role of IT in core object recognition is predicted by the local neuronal discriminability

Finally, with respect to *Prediction 3* of our decoding hypothesis, we found that behavioral deficits from inactivating millimeter scale regions of IT are consistent with predictions from a spatially distributed readout of neurons in IT (Majaj et al. 2015). Indeed, inactivation deficits were well predicted by local neuronal discriminability decoding models, suggesting that the causal role of each IT sub-region is well approximated by the information that is coded explicitly (i.e. linearly separably) by the local population of neurons. In contrast, inactivation deficits were not well predicted by specific local neural response readout models, which predict that neurons that respond highly to particular stimulus classes, without explicitly encoding the differences between them, are causally involved in discrimination between these classes. None of the tested decoding models perfectly explain the inactivation deficits, potentially due to data limits. In the current work, we did not have sufficient neural sampling to directly test population decoding models (e.g. by simulating perturbations on a localized sub-population within a representative sample of all of IT, and measuring the resulting simulated behavior). Nevertheless, our results are consistent with at least one decoding hypothesis (Majaj et al. 2015) (see Figure 5B).

Importantly, our results directly speak to questions of longstanding interest in systems and cognitive neuroscience. A belief held by many in this field is that the overall responsiveness of a cluster of neurons is indicative of its causal role in behavior. For example, one might conclude that face selective regions, which respond preferentially to images of faces, must causally support face detection and discrimination behaviors (Tsao & Livingstone 2008). An alternative hypothesis, used as the basis for techniques such as multivoxel pattern analysis (MVPA) (Norman et al. 2006), is that the behavioral role of a cluster of neurons is not determined by its responsiveness per se, but by its discriminability. To date, there have been a handful of attempts in human cognitive neuroscience to discriminate between these alternative hypotheses using coarse perturbations of neural activity (e.g. trans-cranial magnetic stimulation, and electrical stimulation (Parvizi et al. 2012, Schalk et al. 2017, Pitcher et al. 2007)). In this study, we were able to both record from and inactivate the neuronal activity in arbitrarily selected millimeter-scale regions in primate IT. In contrast to previous human cognitive neuroscience studies, we found that responsiveness is not at all predictive of the behavioral deficits resulting from inactivation. Instead, our results are consistent with a decoding hypothesis based on neuronal discriminability (Majaj et al. 2015), and demonstrate that one should not conclude that a cluster of neurons that preferentially responds to a particular group of stimuli causally supports the ability to discriminate between stimuli within that group.

## Methods

### Subjects and surgery

Two adult male rhesus macaque monkeys (Macaca mulatta, subjects M, P) were trained on the core object recognition paradigm described below. For each animal, a surgery using sterile technique was performed under general anaesthesia to implant a titanium head post to the skull using titanium screws, and a cylindrical recording chamber (19 mm inner diameter; Crist Instruments) over a craniotomy targeting the temporal lobe in the left hemisphere from the top of the skull (Monkey M, +13 mm posterior-anterior, +16.3 mm medial-lateral, 15° medial-lateral angle; Monkey P, +13 mm posterior-anterior, +14.75 mm medial-lateral, 15° medial-lateral angle). All procedures were performed in compliance with the guideline of National Institutes of Health and the American Physiological Society, and approved by the MIT Committee on Animal Care.

### Core object recognition behavioral paradigm

Core object discrimination is defined as the ability to discriminate between two or more objects in visual images presented under high view uncertainty in the central visual field (~ 10°), for durations that approximate the typical primate, free-viewing fixation duration (~ 200 ms) (DiCarlo & Cox 2007, DiCarlo et al. 2012). As in our previous work (Rajalingham et al. 2015, 2018), we investigate this behavior using batteries of trial-by-trial interleaved set of pairwise object discrimination tasks. The behavioral paradigm is described below. Behavioral data was collected under head fixation, and subjects reported their choices using their gaze. We monitored eye position by tracking the position of the pupil using a camera-based system (SR Research Eyelink 1000). Images were presented on a 27” LCD monitor (1920 × 1080 at 60 Hz; Samsung S27A850D) positioned 44 cm in front of the animal. At the start of each training session, subjects performed an eye-tracking calibration task by saccading to a range of spatial targets and maintaining fixation for 800ms. Calibration was repeated if drift was noticed over the course of the session.

Figure 1B illustrates the behavioral paradigm. Each trial was initiated when the monkey acquired and held gaze fixation on a central fixation point for 200ms, after which a test image (8 × 8° of visual angle in size) appeared at the center of gaze for 100ms. Trials were aborted if gaze was not held within ±2°. After extinction of the test image, two choice images, each displaying a single object in a canonical view with no background, were immediately shown to the left and right (each centered at 8^°^ of eccentricity along the horizontal meridian; see Fig. 1B). One of these two objects was always the same as the object that generated the test image (i.e. the correct choice), and its location (left or right) was randomly chosen on each trial. The object that was not displayed in the test image is referred to as the distractor object, but note that objects are equally likely to be distractors and targets. The monkey was allowed to freely view the choice images for up to 1000ms, and indicated its final choice by holding fixation over the selected image for 700ms. The monkey was rewarded with a small juice reward for each correct trial. After the end of each trial, another fixation point appeared, cueing the next trial. Each trial consisted of a different randomly selected pairwise object discrimination task. Note that each pairwise task is operationally defined by the pair of choice objects at the end of the trial, and we insure that the test images are chosen in a balanced way such that approximately half of the trials begin with test images of one object and the other half of the trials begin with test images of the other object. Performance of each such “pairwise task” is the primary unit of measure in this study (averaged over all test images of each object, unless otherwise noted). Note that, because the trials of each such pairwise discrimination task are randomly interleaved, the subject cannot anticipate which object will be shown or which pair of object choices will appear after the test image. Real-time experiments for monkey psychophysics were controlled by open-source software (MWorks Project http://mworks-project.org/).

Both animals were previously trained on other images of other objects, and were proficient in discriminating among over 35 object categories (i.e. several hundreds of possible pairwise object discrimination tasks). In this study, five basic-level objects were tested (bear, elephant, dog, airplane, and chair) that were picked at random. While other (unpublished) work suggests that more objects are needed to fully exercise the domain of core object recognition, in this study our primary goal was to balance between spanning that domain and collecting enough behavioral trails to detect even subtle changes in discrimination performance that might result from suppression of IT sub-regions. Our choice of five objects resulted in ten possible pairwise object discrimination tasks (see Figure 1A for complete list). To accumulate enough trials to precisely measure performance for each task within a single behavioral session (i.e. a single experimental day), we sub-selected six of these ten tasks for most experiments. For a subset of experiments in one animal (monkey P, experiment 2), we tested all ten pairwise tasks. For each session, monkeys were tested for several hours (until satiation) and performed a large number of trials (monkey M: 3442±1097, monkey P: 4430±942; mean ± SD).

### Test images

We examined basic-level object recognition behavior by generating test images of the five objects (above) that were synthesized from the five computer models of each object. As in prior work (Rajalingham et al. 2015, Majaj et al. 2015), the goal was to use naturalistic images that also exercised the view invariance challenges of core object recognition, and the image generation pipeline is described in detail elsewhere (Majaj et al. 2015). Briefly, each image was generated by first rendering the object with randomly chosen viewing parameters (2D position, 3D rotation and viewing distance), and then placing that foreground object view onto a randomly chosen, natural image. Object models spanned basic-level object categories (bear, elephant, dog, airplane, and chair). Background images were sampled randomly from a large database of high-dynamic range images of indoor and outdoor scenes obtained from Dosch Design (www.doschdesign.com). This image generation procedure enforces invariant object recognition as it requires the animal to tackle the invariance problem, the computational crux of object recognition (Ullman & Humphreys 1996, Pinto et al. 2008). Note that this design is in contrast to many prior perturbation studies of IT cortex in which the subject is required only to match one *image* to that same image (a.k.a. standard “match-to-sample”) (Horel et al. 1987, Biederman et al. 1997), while here the subject must match *any possible image* of an object to a visual token (canonical view) that stands for that object.

The majority of the behavioral data presented here were collected in response to a base image set generated from the five objects (40 test images of each object, 200 test images in total). We additionally generated a variant of this dataset consisting of texture-less images of the same objects. These texture-less images were targeted to both titrate the task difficulty and further remove potential low-level confounds (e.g. luminance and contrast). This texture-less image set was not held fixed in size: on each behavioral session, we tested subjects on a mixture of 20% previously seen and 80% completely novel texture-less images of the same five objects, to mitigate potential memorization strategies. For the purpose of the current work, we treat both of these image sets as equivalent, namely as images of the same five objects under study differing only in their precise generative parameters. Figure S1 shows example two images for each object, from both image sets.

### Physiology and pharmacology

In each animal, we first recorded multi-unit activity (MUA) from randomly sampled sites on the ventral surface of IT (monkey M: 57 multi-unit sites, monkey P: 43 multi-unit sites). Recordings in each animal were made over a period of several weeks using glass-coated tungsten micro-electrodes (impedance, 0.3 – 0.5*M*Ω; outer diameter, 310um; Alpha Omega). A motorized micro-drive (Alpha Omega) was used to lower electrodes through a 26-gauge stainless-steel guide tube inserted into the brain (5 mm) and held by a plastic grid inside the recording chamber (CRIST). We recorded MUA responses from IT while monkeys passively fixated images in a rapid serial visual presentation (RSVP) protocol (10 images/trial, 100ms on, 100 ms off). To ensure accurate stimulus presentation, eye position was tracked and trials were aborted if gaze was not held within ±1.5°. To ensure accurate stimulus locking, spikes were aligned to a photodiode trigger attached to the display screen. Multi-unit responses were amplified (1x head-stage), filtered (250Hz cutoff), digitized (sampling rate of 40kHz) and sorted (Plexon MAP system, Plexon Inc.). For each image and multi-unit site, the image response patterns were obtained by first averaging MUA over many (~ 10) image repetitions, and computing the number of repetition-averaged spikes in two post-stimulus windows (70-170ms, 170-270ms)

Following this mapping stage, we performed inactivation experiments using focal microinjections of muscimol, a potent GABA agonist (Andrews & Johnston 1979). We varied the location of microinjections to randomly sample the ventral surface of IT (from approximately +8mm AP to approx +20mm AP). Given the relatively long half-life of muscimol, inactivation sessions were interleaved over days with control behavioral sessions. Thus, each inactivation *experiment* consisted of three behavioral sessions: the baseline or pre-control session (1 day prior to injection), the inactivation session, and the recovery or post-control session (2 days after injection). Each inactivation session began with a single focal microinjection of 1*μ*1 of muscimol (5mg/ml, Sigma Aldrich) at a slow rate (100nl/min) via a 30-gauge stainless-steel cannula at the targeted site in ventral IT. Injections were made through a simple microinjection circuit consisting of a three-way valve (Labsmith) and marker line (similar to (Noudoost & Moore 2011)), enabling precise monitoring of the flow and volume of muscimol injected. In pilot experiments, we verified complete neural suppression at the location of injection using custom-built single-use injectrodes (Noudoost & Moore 2011). Given this volume of muscimol, we estimate strong neural suppression within a local region of ~ 2.5mm in diameter for up to six hours after injection (Arikan et al. 2002). After completion of the injection, we waited 10-20 minutes before measuring the monkey’s behavior on a battery of object recognition tasks for up to 3 hours post-injection.

To ensure accurate targeting of IT and reconstruction of the relative positions of injection and recording locations, all electrophysiological recordings and pharmacological injections were made under micro-focal stereo x-ray guidance (Cox et al. 2008). Briefly, monkeys were fitted with a plastic frame (3 × 4 cm) positioned near the temporal lobe using a plastic arm anchored in the dental acrylic implant. The frame contained six brass fiducial markers (1mm diameter) of known geometry, measured using micro-CT. The fiducial markers formed a fixed 3D skull-based coordinate system for registering all physiological recordings and pharmacological injection sites. At each site, two x-rays were taken simultaneously at near orthogonal angles, and the 3D location of the electrode/cannula tip was reconstructed relative to the skull using stereo-photogrammetric techniques. This procedure enables high-resolution reconstruction (<200um error) of electrode and cannula locations across experimental sessions (Cox et al. 2008, Issa et al. 2013). Under assumptions of approximate planarity for the ventral surface of IT, we measured the distance between sites in IT using the Euclidean distance between x-ray reconstructed 3-D coordinates.

In total, we collected data for 25 inactivation experiments, with each inactivation experiment consisting of three consecutive behavioral sessions each, in two monkeys (monkey M: *n* = 10 experiments, monkey P: *n* = 15 experiments). Interleaved within this inactivation data collection, we additionally collected behavioral data for 18 control experiments, where each experiment again consisted of three consecutive control behavioral sessions each, with the same images and tasks but with no injections, in both monkeys (monkey M: *n* = 5 experiments, monkey P: *n* = 13 experiments). These control data were used to estimate the natural variability in performance across behavioral sessions.

### Analysis

#### Behavioral metrics

We previously introduced several metrics to characterize behavior in this pairwise object discrimination paradigm (Rajalingham et al. 2018). Here, we focus on the highest resolution behavioral metric that can be reliably measured in a single behavioral session, the one-versus-other object level performance metric (previously termed B.O2). Briefly, this metric is a pattern of pairwise object discrimination performances. For each pairwise object discrimination task, performance was estimated using a sensitivity index *d*’ (Macmillan 1993): *d*’ = *Z*(hit rate) – *Z*(false alarm rate), where *Z*(.) is the inverse of the cumulative Gaussian distribution. All *d*’ estimates were constrained to a range of [0,5].

Recall that each inactivation experiment consisted of three behavioral sessions. We first equated the number of trials per session by selecting the first *N* trials of each session, where *N* was the minimum number of trials across the three sessions. For each of these three behavioral sessions, we then computed a pattern of performances across tasks. We defined the control behavioral performance as the average of the pre-control and post-control performances: *ψ_control_* = (*ψ_precontrol_* + *ψ_postcontrol_*)/2. To measure the behavioral deficit from inactivation, we estimated a behavioral deficit pattern (*δ*) as the difference between inactivated and control performance over tasks:

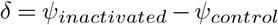

We additionally estimated a normalized behavioral deficit pattern as

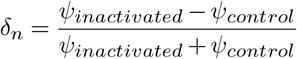

#### Sparsity of deficit

We quantified the non-uniformity of the behavioral deficits using a sparsity index *SI*(*x*) (Vinje & Gallant 2000) as follows:

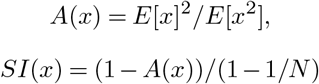

where *E*[.] denotes the expectation of, and *N* is the length of the vector *x*. When applied to a behavioral deficit pattern with no sampling noise, *SI*(*δ*), this index has a value of 0 for perfectly uniform deficit patterns, and a value of 1 for perfectly one-hot deficit pattern. To ensure that the sparsity of the behavioral deficit did not purely reflect non-uniformity in the behavioral difficulty across tasks, we additionally computed this index from the normalized deficit pattern vector *SI* (*δ_n_*). We computed the *SI* for each inactivation site, and estimated the average across all sites.

To ground this empirical *SI* value in intuition, we estimated the corresponding SI distributions for different simulated behavioral deficit patterns with varying degrees of nonuniformity across tasks, and with comparable sampling noise to that in our actual behavioral data (i.e. a finite number of trials). To estimate the expected SI distribution from a deficit with *P*% of tasks affected, we performed the following simulation. For each inactivation site, we computed an estimate of the deficit pattern (*δ*) from a random bootstrap sample of trials. From this deficit pattern estimate, we set all but the top *P*% of deficit values to zero, and randomly shuffled the position of remaining non-zero entries. We averaged the resulting deficit pattern estimates across bootstrap samples to obtain a simulated deficit pattern with approximately equal, non-zero deficit on *P*% of tasks. Finally, we computed the sparsity index for this simulated mean deficit pattern. By varying *P*(= 10%, 25%, …, 100%), we obtained estimates of SI distributions expected from different degrees of non-uniformity across tasks.

#### Neuronal readout models

To investigate the link between neuronal activity and behavioral deficits, we constructed and tested a number of decoding models. Each of these models predicts an inactivation pattern from the activity of neurons recorded in close anatomical proximity (within 2mm) to the injection site. As described above, multi-unit neuronal activity was measured in response to the same images under a passive viewing paradigm and could thus be used as the input to each decoding model. We constructed a feature matrix *R* from the firing rate responses over images (averaged over repetitions) all local multi-unit sites. Each tested decoder model maps *R* to a behavioral deficit prediction Δ. The local neural discriminability and local population discriminability models we tested here were loosely inspired from population readout models of IT (Majaj et al. 2015). Note, however, that the current implementations do not include the remaining (nonlocal) IT population as inputs, as we did not have access to a larger sample of IT. The specific local decoder models we tester here were: *local neural response models* (M1, M2) predict largest deficits for tasks with images that yielded largest response from the local neuronal sites, and *local neural discriminability models* (M3, M4, M5) predict largest deficits for tasks for which the local neural population was most discriminative, as measured by a linear classifier. The details of these five models are as follows:

1. M1 (mean neural response): The deficit for each task Δ_*i,j*_ is estimated as the (negative of) neural response to objects *i,j*, averaged over sites and images ({*R*}_*sites,images*_).
2. M2 (weighted mean neural response): The deficit for each task Δ_*i,j*_ is estimated as the (negative of) neural response to objects *i,j,* averaged over sites and images after weighting each site by its overall discriminability *W* ({*wR*}_*sites,images*_).
3. M3 (local neural response discriminability): The neural image responses averaged over sites, ({*R*}_*sites*_), is used as a single neural feature *f* to train and test a linear SVM. The deficit for each task Δ_*i,j*_ is estimated as the (negative of) the SVM performance (in units of *d*’) to objects *i, j*, averaged over images.
4. M4 (local neural discriminability, mass action): Each neural site’s image response (*R_site_k__*), is used as a single neural feature *f* to train and test a linear SVM. The deficit for each task Δ_*i,j*_ is estimated as the (negative of) the SVM performance (in units of *d*’) to objects *i,j*, averaged over images, summed over all sites *k.*
5. M5 (local population discriminability): The local neuronal image response (*R*), is used to train and test a linear SVM. The deficit for each task Δ_*i,j*_ is estimated as the (negative of) the SVM performance (in units of *d*’) to objects *i,j,* averaged over images.

#### Noise-adjusted correlations

We measured the similarity between two behavioral deficit patterns *δ*_1_, *δ*_2_ (e.g. between true deficit patterns and predictions from a model) using a noise-adjusted correlation (DiCarlo & Johnson 1999, Johnson et al. 2002). For each behavioral deficit pattern, we split all independent raw observations (e.g. behavioral trials) into two equal halves and computed the behavioral deficit pattern from each half, resulting in two independent estimates of the deficit pattern. We took the Pearson correlation between these two estimates as a measure of the reliability of that behavioral deficit pattern, given the data, i.e. the split-half internal reliability. To estimate the noise-adjusted correlation between two deficit patterns, we compute the Pearson correlation over all the independent estimates of deficits from each, and we then divide that raw Pearson correlation by the geometric mean of the split-half internal reliability of each deficit:

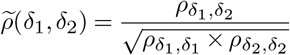

Since all correlations in the numerator and denominator were computed using the same amount of trial data (exactly half of the trial data), we did not need to make use of any prediction formulas (e.g. extrapolation to larger number of trials using Spearman-Brown prediction formula). This procedure was repeated 10 times with different random split-halves of trials. Our rationale for using a reliability-adjusted correlation measure was to account for variance in the behavioral deficit that is not replicable by the task condition. If two behavioral deficits are identical, then their expected noise-adjusted correlation is 1.0, regardless of the finite amount of data that are collected. The noise-adjusted correlation was used to compute the similarity between observed and predicted behavioral deficit patterns (e.g. for testing neural readout models), as well as for the similarity between two different behavioral deficit patterns arising from two different inactivation sites.

#### Statistical testing

Unless otherwise specified, we estimated the uncertainty in behavioral deficit measurements (i.e. delta, see above) via bootstrap resampling of trials, repeated 100 times. The standard error of each delta measurement was estimated as the standard deviation of its bootstrap distribution. For statistical tests, we performed one-tailed exact tests, by computing the empirical probability of observing a sample below zero. To compute this probability from the empirical bootstrap distribution, we fit a Gaussian kernel density function to the empirical distribution, optimizing the bandwidth parameter to minimize the mean squared error. This kernel density function was evaluated to compute a p-value, by computing the cumulative probability of observing a positive behavioral delta.

## ACKNOWLEDGEMENTS

We thank Arash Afraz, Nancy Kanwisher and Roozbeh Kiani for useful conversations. This research was supported by US National Eye Institute grants R01-EY014970 (J.J.D.), Office of Naval Research MURI-114407 (J.J.D), and the Simons Foundation SCgB [325500] (J.J.D.).

## Supplemental Materials

**Fig. S1.**
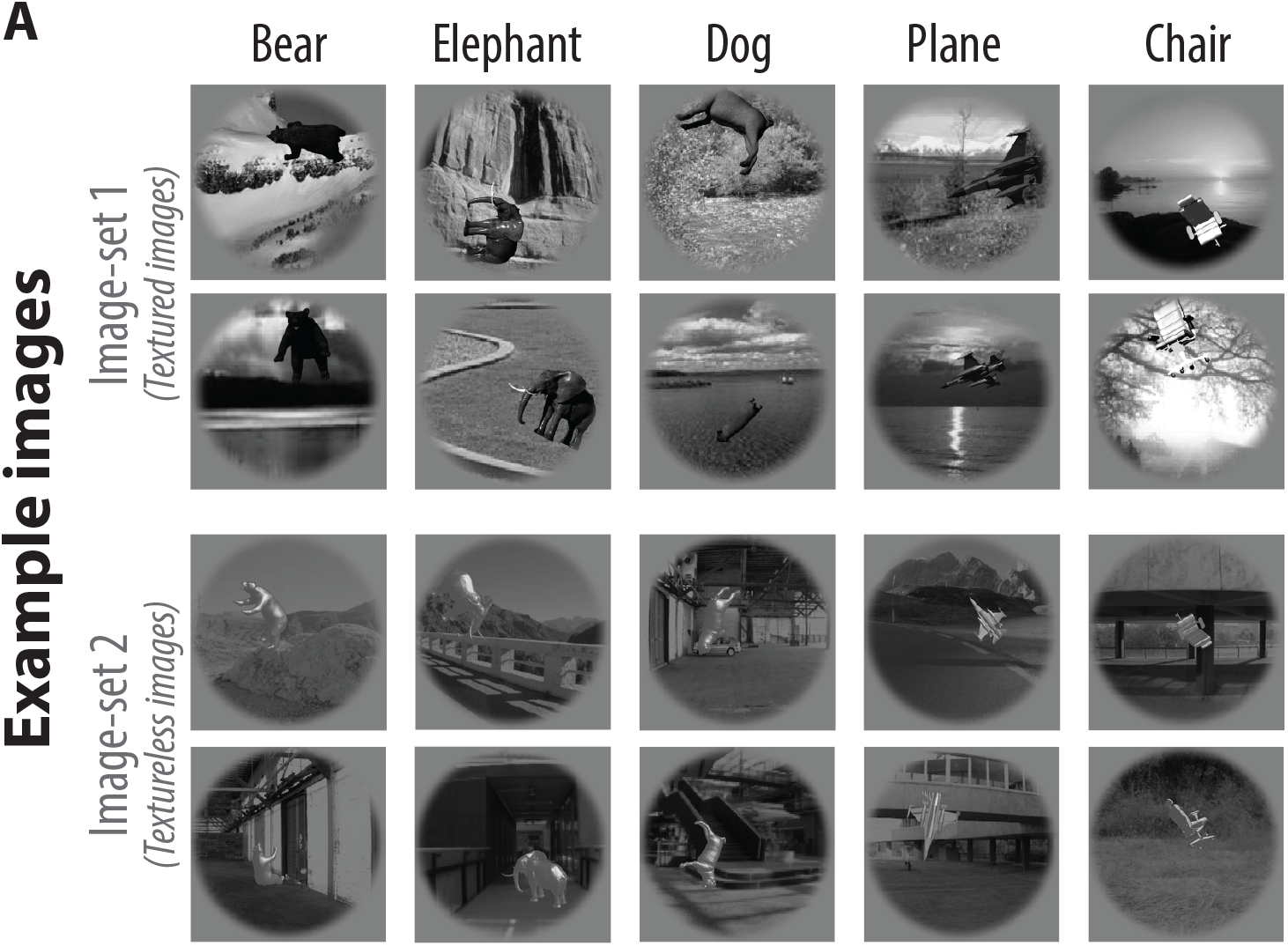
(a) Visual images. Two (out of hundreds) example images per object, for each of the five objects and for both image sets, are shown. Stimuli consisted of naturalistic synthetic images of 3D objects rendered under high view-uncertainty and overlaid on a naturalistic background. We additionally generated a dataset consisting of texture-less images of the same objects; two example texture-less images for each of the five objects are shown (image-set 2). For the purpose of the current work, we treat both of these image sets as equivalent, namely as images of the same five objects under study.

**Fig. S2.**
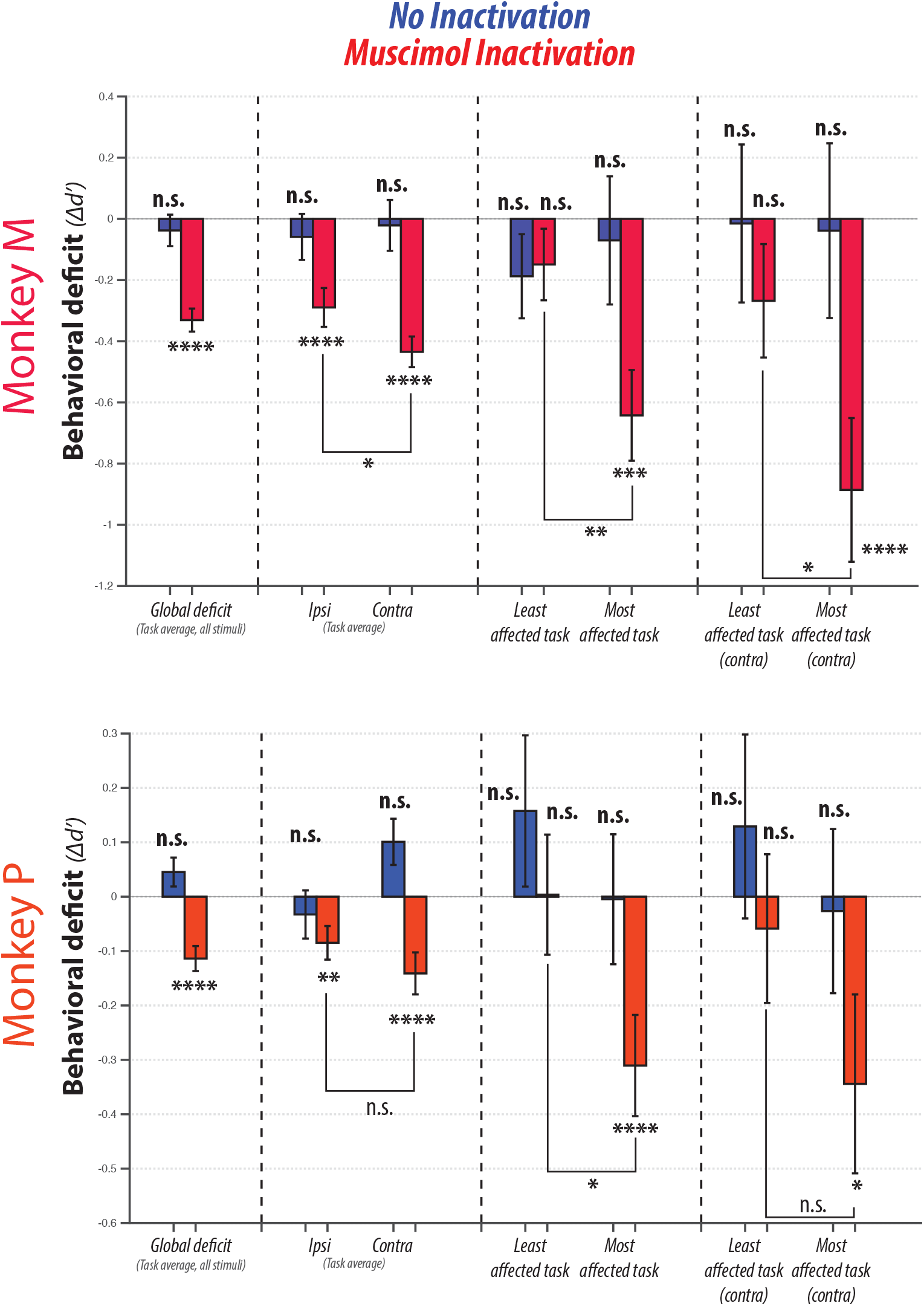
Summary of behavioral deficits for each monkey. Formatting as in Figure 3B

